# Age and Sex Influence Diurnal Memory Oscillations, Circadian Rhythmicity, and *Per1* Expression

**DOI:** 10.1101/2025.07.24.666612

**Authors:** Lauren Bellfy, Gretchen C. Pifer, Megan J. von Abo, Chad W. Smies, Alicia R. Bernhardt, Achintya Perumal, Madison J. Jackson, Janine L. Kwapis

## Abstract

**Background:** The circadian system influences many different biological processes across the lifespan, including memory performance and daily activity patterns. The biological process of aging causes decreased control of the circadian system that is accompanied by a decline in memory performance, suggesting that these two processes may be linked. Indeed, our previous work has shown that in male mice, the clock gene *Per1* functions within the dorsal hippocampus to exert diurnal control over memory and repression of *Per1* in the old hippocampus contributes to age-related impairments in spatial memory. Although it is clear that *Per1* may be a key molecular link between memory and the circadian rhythm, next to nothing is known about how sex impacts this role in the young or old brain. Here, we are interested in understanding how the factors of sex and age impact memory performance, circadian activity patterns, sleep behavior, and hippocampal *Per1* expression.

**Methods:** We used a combination of spatial memory (Object Location Memory (OLM)) and circadian activity monitoring to determine how male and female mice change across the lifespan. In addition, we used RT-qPCR to quantify the change in *Per1* levels in response to learning in young and old, male and female mice.

**Results:** Young female mice resist diurnal oscillations in memory, showing robust spatial memory across the diurnal cycle. In contrast, old female mice show an emergence of diurnal memory oscillations, with better memory during the day than at night (similar to what we observed previously in young male mice). In contrast, old male mice showed better memory performance during the night than the day, suggesting that their peak memory performance is drastically shifted compared to young males. We also measured activity patterns and sleep behavior across the diurnal cycle and found that sex was more of an influence than age in multiple analyses, but age did have an impact, with old male mice showing stronger circadian rhythm disruptions than any other cohort. Finally, we investigated whether the circadian clock gene *Per1* plays a role in these sex- and age-dependent effects in diurnal memory performance. We found that, in general, learning-induced *Per1* and memory performance peaked at similar times of day in each group, consistent with our hypothesis that *Per1* exerts diurnal control over memory performance.

**Conclusions:** This work supports a role for *Per1* in exerting diurnal control over memory and suggests that *Per1* may be an appealing therapeutic target to improve memory and circadian dysfunction in old age.

**Highlights:**

- Diurnal oscillations in spatial memory are sex- and age-dependent in mice
- *Per1* learning-induced expression matches diurnal memory patterns
- Circadian rhythm patterns are sex- and age-dependent in mice
- Young females show good memory across the diurnal cycle
- Diurnal memory oscillations reemerge in old female mice

**Plain language summary:** Memory is an integral part of everyday functioning, and one that is known to decline with aging. Our lab has previously shown that the clock gene *Period1* (*Per1)* regulates spatial memory performance in young males, establishing a molecular link between circadian rhythms and memory. Young adult male mice show diurnal oscillations in memory consolidation, with the best memory occurring at midday, and the worst memory occurring at midnight. In the current study, we wanted to expand our work to young adult females, as well as an aged population of male and female mice. Using a simple spatial memory task, we measured diurnal changes in both memory performance and *Per1* gene expression within the dorsal hippocampus (a brain region necessary for spatial memory). We found that old mice (both male and female) showed a correlation between high *Per1* levels and better memory, as we have previously seen.

Conversely, young female mice performed well on the memory task at every timepoint but didn’t have a significant change in *Per1*, indicating that they may be using some different mechanism to modulate memory performance. Finally, we used infrared activity monitoring to investigate several circadian rhythm related measures in young and old, male and female mice. We found that sex influenced the circadian rhythm more than age, and the group with the largest circadian disruption was aged males. Overall, this research provides new information about how both sex and age impact diurnal oscillations in both memory and activity, fundamental knowledge that has been lacking in the field.

## Introduction

Circadian rhythms regulate myriad physiological functions, including cardiovascular and metabolic functioning, feeding behaviors, sleep-wake timing, activity, body temperature, and cognitive functioning [1]. Previously, we have shown that memory consolidation is one biological function that is tightly regulated by this daily rhythm in young male mice; spatial memory is better during the day than at night in these animals [2]. Further, we found that there is a corresponding increase in the core clock gene *Period1 (Per1)* within the dorsal hippocampus (DH) and retrosplenial cortex that supports spatial memory; *Per1* is only induced by spatial learning during the daytime, when memory is robust, and is not induced at night when memory is poor [2–4]. Manipulating *Per1* expression in either brain region modulates memory, with local *Per1* knockdown impairing daytime memory and overexpression improving memory in old mice with memory impairments [2–4]. This suggests that *Per1* is a key mechanism capable of exerting local diurnal control over memory within multiple memory-relevant brain regions.

*Per1* is best known for its canonical role as one of the core clock genes that oscillates within the suprachiasmatic nucleus (SCN) to regulate the circadian system. The central molecular machinery of the circadian system consists of an autoregulatory transcription-translation negative feedback loop that begins when two proteins, CLOCK and BMAL1, form a heterodimer. The CLOCK-BMAL1 dimer then binds to E-box motifs upstream of the *Per* and *Cry* gene families, driving their transcription. These transcripts move to the cytoplasm for translation and their proteins heterodimerize before translocating back into the nucleus to inhibit subsequent CLOCK/BMAL1 activity. Thus, PER and CRY ultimately inhibit their own transcription, closing the negative feedback loop. Eventually, the PER and CRY protein products are degraded, freeing up the CLOCK/BMAL1 heterodimer to allow a new round of transcription. This whole feedback loop takes approximately 24 hours to complete, which sets circadian rhythm length in individual cells, both within the SCN and in satellite clocks like the dorsal hippocampus [5–8]. Importantly, the length and regularity of this feedback loop changes with aging, typically becoming longer and more disrupted [9–11]. Age-related changes in *Per1* dynamics across a range of satellite clocks could therefore contribute to multiple aspects of the aging process.

Consistent with this idea, work from our lab has suggested that hippocampal *Per1* may play a critical role in age-related memory decline. Normal aging is accompanied by decreased cognition, including impairments in memory, in both humans and rodents [3,4,12–14]. Further, our past work has shown that abnormal epigenetic repression in the old male hippocampus prevents local induction of *Per1* in response to learning, contributing to age-related spatial memory impairments [3]. This suggests that repression of *Per1* in the old hippocampus might contribute to a persistent ‘nighttime state’ that limits memory even during the daytime, when memory is normally robust. However, to date, all of the work on *Per1*’s role in hippocampal memory has focused exclusively on male mice and it is therefore unclear how sex impacts circadian rhythmicity, sleep*, Per1* expression, and/or memory performance in the young and old brain.

Here, we tested how diurnal oscillations in memory and the accompanying *Per1* induction are affected by both age and sex using the DH-dependent object location memory (OLM) task. First, we used OLM to assess how spatial memory performance varies across the diurnal cycle in young and old female mice and aged male mice. We expected that young female mice would look similar to young males [2], showing a diurnal oscillation in memory that peaks during the daytime. We further expected that memory would be consistently disrupted in old age, with old males and females showing little memory regardless of the time of day. Instead, we found that young female mice show no oscillation in memory, with robust memory even at night. Further, we found that old male mice are capable of acquiring robust memory, just at a different time of day, showing an unexpected peak in memory at night. Finally, as female mice age, oscillations in memory emerge, with old female mice showing peak memory during the day and a trough at night. Next, to determine whether *Per1* might contribute to these sex- and age-dependent results, we measured *Per1* mRNA in the DH, and found that, overall, *Per1* induction levels matched memory performance across the age and sex differences, consistent with our hypothesis that *Per1* is capable of modulating memory across the diurnal cycle. Finally, we used infrared activity monitoring to determine if circadian activity patterns, photic phase resetting, or gross sleep behavior differ with sex or age. We found that sex impacted both activity and sleep, with females showing more overall activity and less sleep than males, regardless of age. Across all of the different measures of activity and sleep, old males showed the most severe disruptions, suggesting that males are especially vulnerable to age-related dysregulation of the circadian system. Overall, this work demonstrates two important findings. First, young female mice show an unexpected resistance to diurnal troughs in memory. This is accompanied by a lack of *Per1* induction at any time of day, suggesting that young females may rely on a different mechanism to support robust memory across the diurnal cycle. Second, aging does not produce consistent impairments in memory across the diurnal cycle, as expected, but shifts the peak performance to an unexpected time-of-day, with male mice showing robust memory in the middle of the night. Rather than inducing a persistent nighttime state, aging may therefore shift memory oscillations so they no longer align with the animal’s diurnal needs. Diurnal oscillations in memory and accompanying *Per1* induction are therefore regulated by both sex and age in unanticipated ways.

## Materials and Methods

### Mice

Young (2-4 months) and old (19-22 months) male and female C57BL/6J mice were used for behavioral experiments. In total, 9 young male, 96 young female, 112 old male, and 103 old female mice were used in the final data. Mice were housed with food and water *ad libitum* under a 12 h light/dark cycle, with lights turning on at 6 am (7 am Daylight Saving Time), for all OLM experiments. Mice that were trained during the day (ZT1, 5, and 9) were entrained under a standard light cycle while mice that were trained during the night (ZT13, 17, 21) were entrained to a reverse light cycle (lights off at 6/7am). During the circadian monitoring experiment, mice were housed under a 12 h light/dark cycle (LD) for 7 days before being housed in constant darkness (24 h dark/dark cycle, DD). During this time, animals were monitored and cages changed in the dark using infrared googles to prevent mice from being exposed to any outside light source. During the experiments testing memory performance, mice were group housed (2-4/cage) by sex and age. For RT-qPCR and circadian monitoring experiments, all mice were single housed (to prevent excessive baseline gene expression or to enable infrared circadian activity monitoring, respectively) and received a wood chew bar for enrichment. All experiments were performed according to US National Institutes of Health guidelines for animal care and use and were approved by the Institutional Animal Care and Use Committee of the Pennsylvania State University.

### Object Location Memory (OLM)

OLM was conducted as previously described [2,3,15]. Mice were handled for 2 min/day for 4 days and habituated to the context without objects for 5 min/day for 6 days. For training, mice were exposed to two identical objects (100 mL glass beakers filled with cement) placed in specific locations within the familiar context for 10 minutes. The following day (24h later) mice were tested in the same context for 5 minutes with one of the of the objects moved to a new location. Object exploration in each session was hand-scored by blinded experimenters, and exploration was defined as the mouse orienting itself towards the center of the object when it was within 1 cm of the object, with digging and rearing behavior not included in exploration time [2,3,15]. Total investigation time was quantified using Deepethogram [16] and the discrimination index (DI) was calculated based off the ratio of time spent investigating the novel location versus the familiar location using the equation: DI = ((novel location exploration - familiar location exploration)/total exploration) x 100. Any mouse displaying a DI more than two standard deviations from the mean was removed, as well as mice that failed to explore sufficiently were removed from all analyses using the following exploration thresholds: Young: removed if exploration <2.5s during testing or <4s during training. Old: removed if exploration <2s during testing or <4s during training. For both ages, mice were removed if they demonstrated an object preference during training (DI >±20). All habituation sessions were analyzed utilizing Ethovision (Noldus, Leesburg, VA) for both average speed and total distance traveled during the session. For molecular experiments, OLM was conducted identically through training, except that mice were sacrificed via cervical dislocation exactly 1 hour after acquisition to capture peak gene expression [17]. The brains were harvested and flash-frozen using isopentane.

Habituation, training and testing sessions for all groups were conducted under dim red light to avoid light-evoked induction of clock genes and to ensure equivalent behavioral conditions across groups.

### RT-qPCR

Tissue and RNA extraction from dorsal hippocampus punches was followed by cDNA synthesis and RT-qPCR for *Per1* expression as previously described [2,4,17]. RNA was extracted using the RNeasy Minikit (Qiagen, Hilden, Germany) and cDNA was synthesized using the Hi-Fi cDNA Synthesis Kit (Abcam, Cambridge, UK). If possible, RT-qPCR assays were designed to be intron spanning to reduce genomic DNA amplification. *Per1* target assay and *GAPDH* reference assay (IDT, Newark, NJ) were designed based on the Mus_musculus, GRCm38 build. qPCR was run on the Roche LightCycler 96 with the following cycling conditions: 1 cycle at 95°C for 3 min, 45 cycles of 95°C for 15 sec and 60°C for 60 sec, and hold at 4°C. Data were analyzed with LightCycler Analysis software using the Relative Quantification analysis method. For *Per1* we used the following primers: left 5’-CCTGGAGGAATTGGAGCATATC-3’; right 5’-CCTGCCTGCTCCGAAATATAG-3’; probe 5’-AAACCAGGACACCTTCTCTGTGGC-3’ labeled with FAM. For *Gapdh* we used the following primers: left 5’-GGAGAAACCTGCCAAGTATGA-3’; right 5’-TCCTCAGTGTAGCCCAAGA-3’; probe 5’-TCAAGAAGGTGGTGAAGCAGGCAT-3’ labeled with HEX.

### Circadian Rhythm and Sleep Time Analysis

Mice were single housed under a 12 h light/dark (LD) cycle. Activity was continuously monitored with passive infrared sensors and Clocklab software (Actimetrics, Layfette, IN). After 7 days of monitoring, lights were turned off and mice were monitored under constant darkness for 10 days. On day 9, activity data was obtained to calculate the free-running Tau value for each mouse using Clocklab Analysis Version 6 (Actimetrics, Layfette, IN). From the free-running Tau, the circadian time (CT) was calculated for day 10. On day 10, mice were exposed to a 50 lux light pulse for 30 minutes starting at CT16.75 until CT17.25 to initiate light-induced phase resetting. Activity was monitored for an additional 10 days [18] to determine whether this light stimulation initiated a phase shift. Clocklab Analysis Version 6 was utilized for activity onset, Tau, circadian clock resetting, and bout analysis. Sleep behavior was assessed using a program adapted from the COMPASS system [19]. Movement data were collected in 10s bins with the criteria that periods of inactivity lasting 40s or longer were quantified as a behavioral correlate of sleep. Previous work has demonstrated a high degree of correlation between this immobility-defined sleep and sleep defined using EEG records [19]. This sleep behavior (in seconds) was collapsed into 30-minute bins across the day/night cycle. Mice were removed from analyses if they died prior to the end of the experiment or were removed from the light pulse analysis if the pulse was delivered >1 CT from CT16.25-17.25.

### Statistical Analysis

Differences in memory performance were analyzed using one-sample t-tests (to compare each group’s object preference to zero), unpaired t-tests, or one-, two-, or three-way ANOVAs followed by Sidak’s multiple comparison post-hoc tests. For RT-qPCR, each group was normalized to the ZT17 homecage condition (to compare across groups) or to its own time-matched homecage group (to compare the magnitude of learning-induced increases across groups). Sleep behavior and habituation movement data were analyzed with mixed-model ANOVAs with sleep behavior or movement treated as a repeated measures variable. For the various light pulse analyses, the data were analyzed using one sample t-tests, unpaired t-tests, or two- or three-way ANOVAs followed by Sidak’s multiple comparison post-hoc tests. For all analyses, significance was indicated by an α value of 0.05.

## Results

### Memory performance oscillates across the diurnal cycle in old but not young female mice

Our previous work has shown that memory performance oscillates across the diurnal cycle in young (2-4 months old) male mice, peaking during the daytime [2], but it is unknown if young female mice show identical memory oscillations. To test this, we trained young (2-4 months old) female mice in the dorsal hippocampus (DH)-dependent Object Location Memory (OLM) task at six distinct timepoints across the diurnal cycle (Fig. 1A; ZT1, ZT5, ZT9, ZT13, ZT15, ZT17, ZT21, where ZT=lights on, 6/7am). We tested the mice 24 hours following training to assess long-term memory (Fig. 1B). Unlike in our previous work, where young male mice showed significantly worse memory during the night [2], young female mice showed no diurnal oscillation in memory, instead showing robust memory performance across the day/night cycle (Fig. 1C). Animals showed intact memory for OLM at each timepoint except ZT1 (one-sample *t-*tests comparing each group to 0, ZT1: *t*_(7)_=1.210, p>0.05, ZT5: *t*_(8)_=4.304, p<0.01, ZT9: *t*_(7)_=6.361, p<0.001, ZT13: *t*_(9)_=2.819, p<0.05, ZT17: *t*_(7)_=2.062, p>0.05, ZT21: *t*_(13)_=4.312, p<0.001) and there was no significant effect of time-of-day (one-way ANOVA: *F*_(5,51)_=0.9630, p=0.2801) (Fig. 1C). Even when we collapsed across the day and night timepoints, we failed to find a significant difference in memory performance (Fig. 1D; unpaired *t*-test, *t*_(55)_=0.3824, p=0.7036). Finally, there was no significant difference in total object exploration at test across the diurnal cycle (Suppl. Fig. 1A; one-way ANOVA: *F*_(5, 51)_=0.9342, p=0.4668) and no difference in habituation movement during the day or night (Suppl. Fig. 2A; distance traveled: two-way ANOVA: significant effect of habituation day (*F*_(5,330)_=2.318, p<0.05), but no day/night effect or interaction; velocity: two-way ANOVA: significant effect of habituation day (*F*_(5,330)_=2.647, p<0.05), but no effect of day/night or interaction). Interestingly, while the young female mice do show more exploration at night during training (data not shown; one-way ANOVA: *F*_(5, 51)_=4.362, p<0.01), this does not seem to impact the strength of the memory formed, as all groups show similar memory strength at test. These results suggest that young female mice do not show diurnal oscillations in memory and resist the ebbs in memory at night seen in young male mice.

**Figure 1.**
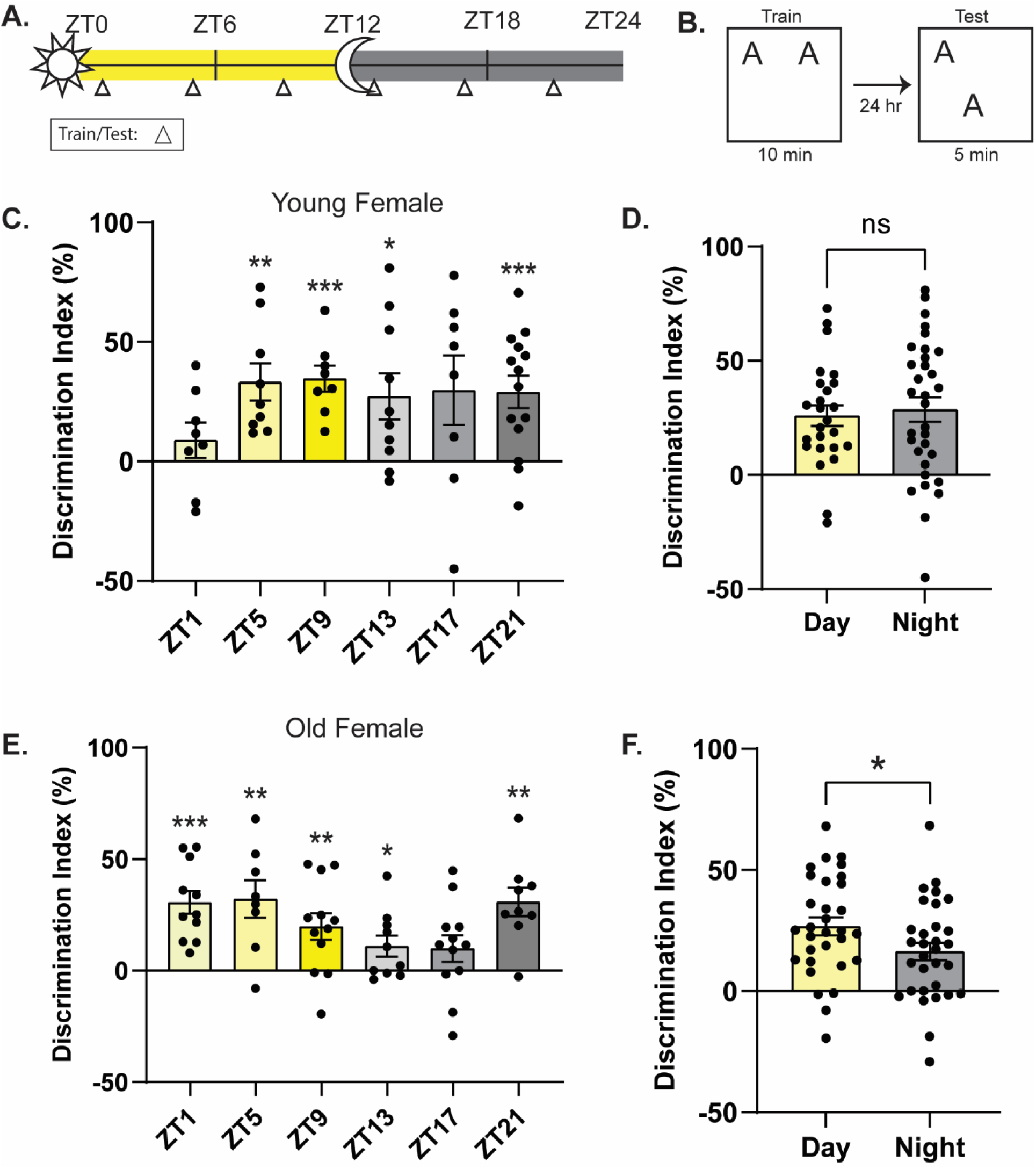
Memory oscillates in old but not young female mice across the diurnal cycle**. A.** Schematic for behavioral timepoints. Triangles indicate time of training and testing. **B.** Object location memory experimental design. **C.** No oscillatory patterns for memory performance are present in young female mice across the diurnal cycle (n=8-14/timepoint). **D.** No difference in memory performance during the day versus the night in young female mice (n=25-32/timepoint). **E.** Memory oscillates across the diurnal cycle in old female mice with the peak of memory at ZT5 and the trough at ZT17 (n=8-12/timepoint). **F.** Memory performance is significantly better during the day than the night in old female mice (n=31/timepoint). * = p<0.05, ** = p<0.01, *** = p<0.001, **** = p<0.0001 compared to zero (**C., E.**) or between groups (**D., F.**). ZT = Zeitgeber Time, where ZT0 = 6am (7am DST), lights on, ZT12 = 6pm (7pm DST), lights off.

Next, we tested how this diurnal regulation of memory might change in old female mice. As aging typically decreases cognitive function across species [12,14,20,21] and previous research has shown that old male mice have a deficit in spatial memory during the daytime, when memory is usually best [2–4,14], we hypothesized that old (19-22 months old) female mice would show weak memory across the diurnal cycle. As before, we trained old female mice on OLM at the same six timepoints and tested each group 24h later (Fig. 1A and 1B). We found that old female mice showed better memory during the day timepoints (ZT1, ZT5, ZT9) than the night timepoints (ZT13, ZT17, ZT21; Fig. 1E; one-way ANOVA: *F*_(5,56)_=2.800, p<0.05; no significant difference between timepoints). Interestingly, as seen with our young male mice, the peak of memory in our old females was at ZT5 and the trough at ZT17 [2] (one-sample *t-*tests comparing each group to 0, ZT1: *t*_(10)_=5.881, p<0.001, ZT5: *t*_(7)_=3.808, p<0.01, ZT9: *t*_(11)_=3.314, p<0.01, ZT13: *t*_(9)_=2.323, p<0.05, ZT17: *t*_(11)_=1.648, p=0.1276, ZT21: *t*_(8)_=4.851, p<0.01). We then compared the collapsed day and night timepoints and found significantly better memory during the day than the night (Fig. 1F; unpaired *t*-test, *t*_(60)_=2.019, p<0.05). Notably, memory performance starts to recover at the ZT21 timepoint, which brings up the average of the night timepoints, although the comparison between day and night was significant. During training there was no difference in exploration or training discrimination index at any timepoint, indicating that this was not a factor in the time-of-day memory performance (not shown). Finally, we found no difference in total object exploration at test across the diurnal cycle (Suppl. Fig. 1B; one-way ANOVA: *F*_(5,56)_=1.190, p=0.3256), indicating exploration time does not impact memory performance. We did observe that nighttime mice showed more movement during habituation (Suppl. Fig. 2C-D; distance traveled: two-way ANOVA: significant effect of habituation day (*F*_(5,360)_=2.358, p<0.05), significant effect of day/night (*F*_(1,360)_=4.680, p<0.05) but no significant interaction; velocity: two-way ANOVA: significant effect of habituation day (*F*_(5,360)_=2.467, p<0.05), significant effect of day/night (*F*_(1,360)_=5.141, p<0.05) but no significant interaction), indicating that these old female mice show slightly more movement at night, when memory is worst. Notably, general movement is reduced during the daytime, when memory is strongest, suggesting that even when movement is lower, the mice are exploring sufficiently to learn the task.

Therefore, contrary to our hypothesis, when female mice age, they show an emergence of diurnal memory oscillations, shifting from robust and non-oscillating memory as young females to having a strong diurnal memory oscillation as old females, with better memory observed during the daytime than at night.

### Memory performance oscillates across the diurnal cycle in old male mice, peaking at night rather than during the day

We were surprised to find that, rather than showing consistent memory impairments across the diurnal cycle, old female mice are capable of forming robust memory at some, but not all times of day. Next, we asked whether this is also true in old male mice. Our previous work demonstrated that old males show impaired memory during the daytime [2–4], but whether they are consistently impaired across the diurnal cycle is unclear. Therefore, we next investigated whether old (19-22 months old) male mice show intact memory performance for OLM at any point in the diurnal cycle. Here we trained and tested old male mice in OLM at the same six timepoints to assess diurnal variations in long-term memory (Fig. 2A and 2B). We found that old males do show a diurnal oscillation in memory performance, with significantly better memory for OLM during the night than the day (Fig. 2C-D; C: one-way ANOVA: *F*_(5,71)_=6.139, p<0.0001, Sidak’s *post-hoc* tests: ZT17 significantly higher than ZT1, p<0.0001, ZT5, p<0.01, ZT9, p<0.05, ZT13, p<0.05, no other comparisons significant; D: collapsed day vs night: Unpaired *t*-test, *t*_(75)_=3.711, p<0.001). Memory peaked at ZT17 and showed a trough at ZT1. Interestingly, our old male mice showed evidence of spatial memory at all timepoints (one-sample *t*-tests comparing each group to 0, ZT1: *t*_(14)_=2.619, p<0.05, ZT5: *t*_(11)_=2.544, p<0.05, ZT9: *t*_(11)_=4.594, p<0.001, ZT13: *t*_(12)_=5.612, p<0.001, ZT17:

**Figure 2.**
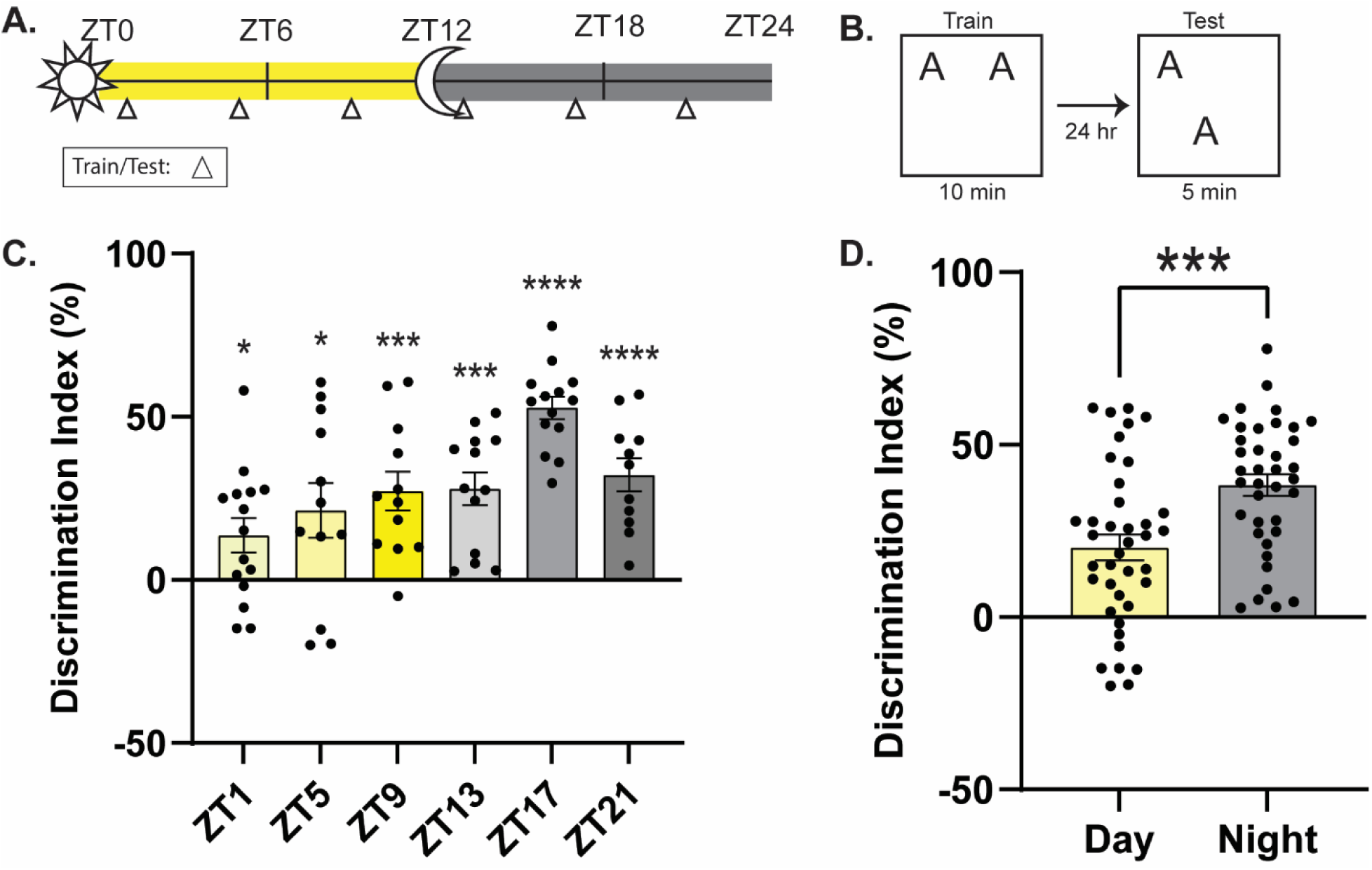
Memory oscillates in old male mice across the diurnal cycle. **A.** Schematic for behavioral timepoints. Triangles indicate time of training and testing. **B.** Object location memory experimental design. **C.** Memory oscillates across the diurnal cycle in old male mice with the peak of memory at ZT17 and the trough at ZT1 (n=10-14/timepoint). **D.** Memory performance is significantly better during the night than the day in old male mice (n=38-39/timepoint). * = p<0.05, *** = p<0.001, **** = p<0.0001 compared to zero (**C.**) or between groups (**D.**). ZT = Zeitgeber Time, where ZT0 = 6am (7am DST), lights on, ZT12 = 6pm (7pm DST), lights off.

*t*_(13)_=15.56, p<0.0001, ZT21: *t*_(10)_=6.305, p<0.0001), while we typically see little to no long-term memory for OLM during the day for mice of this age [3]. Nonetheless, memory performance was weaker during the day than at night (Fig. 2D), consistent with our past work that aging impairs memory during the daytime. We found no significant differences in total object exploration at test (Suppl. Fig. 1C; one-way ANOVA: *F*_(5,71)_=1.172, p=0.3316) and no difference in habituation movement (Suppl. Fig. 2E-F; distance: two-way ANOVA: significant effect of habituation day (*F*_(5,450)_=7.341, p<0.01), but no effect of day/night or interaction; velocity: two-way ANOVA: no effect of habituation day, day/night, or interaction), indicating that memory performance is not dependent on exploration or activity. During training, the old male mice do not show a difference in exploration time or discrimination index at any timepoint, indicating training performance does not drive memory performance at test. Interestingly, these results suggest that aging does not consistently impair memory across the diurnal cycle in old males, as hypothesized, but instead shifts when memory peaks and troughs.

### *Per1* expression is increased in response to learning during the day but not the night in young male and old female mice

Our previous work identified the clock gene *Period1* (*Per1*) as a diurnal regulator of memory consolidation in the hippocampus and retrosplenial cortex of young male mice [2]. We found that *Per1* induction occurs in tandem with memory performance in young males in both regions, peaking when memory is best during the day and showing a trough when memory is worst at night [2,4]. Further, this daytime *Per1* induction is impaired in old mice with age-related memory impairments, consistent with the idea that repressed *Per1* impairs memory. Finally, we showed that manipulating *Per1* impacts memory; knocking down *Per1* in the DH or RSC impairs memory in young males and upregulating *Per1* in these regions ameliorates age-related memory impairments in old males [3]. *Per1* therefore seems to exert diurnal control over memory and daytime repression of *Per1* contributes to the observed deficits in memory during the day in old male mice. To determine if *Per1* plays a similar role in regulating memory across age and sex, we next measured *Per1* levels in each age and sex group at ZT5 or ZT17 (the ‘middle’ of the day or night and when memory peaks/troughs in young males).

Here, we ran male and female mice of each age through 10-min OLM training at ZT5 or ZT17 and sacrificed them along with time-matched homecage control animals 60 minutes later (when learning-induced *Per1* peaks ([17]; Fig. 3A). We found that the time of learning (ZT Time), sex of the animal, and age of the animal all influenced learning-induced *Per1* levels (Fig. 3B; three-way ANOVA, significant effect of ZT time (*F*_(1,51)_=13.74, p<0.001), sex (*F*_(1,51)_=22.91, p<0.0001), ZT time x age (*F*_(1,51)_=4.961, p<0.05), age x sex (*F*_(1,51)_=4.130, p<0.05), ZT time x age x sex (*F*_(1,51)_=19.41, p<0.0001), no significant effect of age and ZT time x sex). As we have previously observed, young male mice showed a significant induction of *Per1* during the daytime that is abolished at night, corresponding to memory performance (also better during the day than at night). Old female mice, which show a similar oscillation in memory to young males ([2], with better memory during the day and worse memory at night) also had a significantly bigger learning-induced increase in *Per1* during the day than at night (Sidak’s *post-hocs* comparing day to night: young male: p<0.0001; old female: p<0.05). In both of these groups, *Per1* expression was significantly induced in trained animals during the day but not at night (Fig. 3B; one sample *t-*tests comparing each group to 100, young male: day: *t*_(5)_=10.11, p<0.001, night: *t*_(7)_=2.210, p=0.0628; old female: day: *t*_(7)_=6.070, p<0.001, night: *t*_(8)_=1.072, p=0.3148). Therefore, young male and old female mice show similar diurnal oscillations in learning-induced *Per1* that correspond to the observed oscillations in memory.

**Figure 3.**
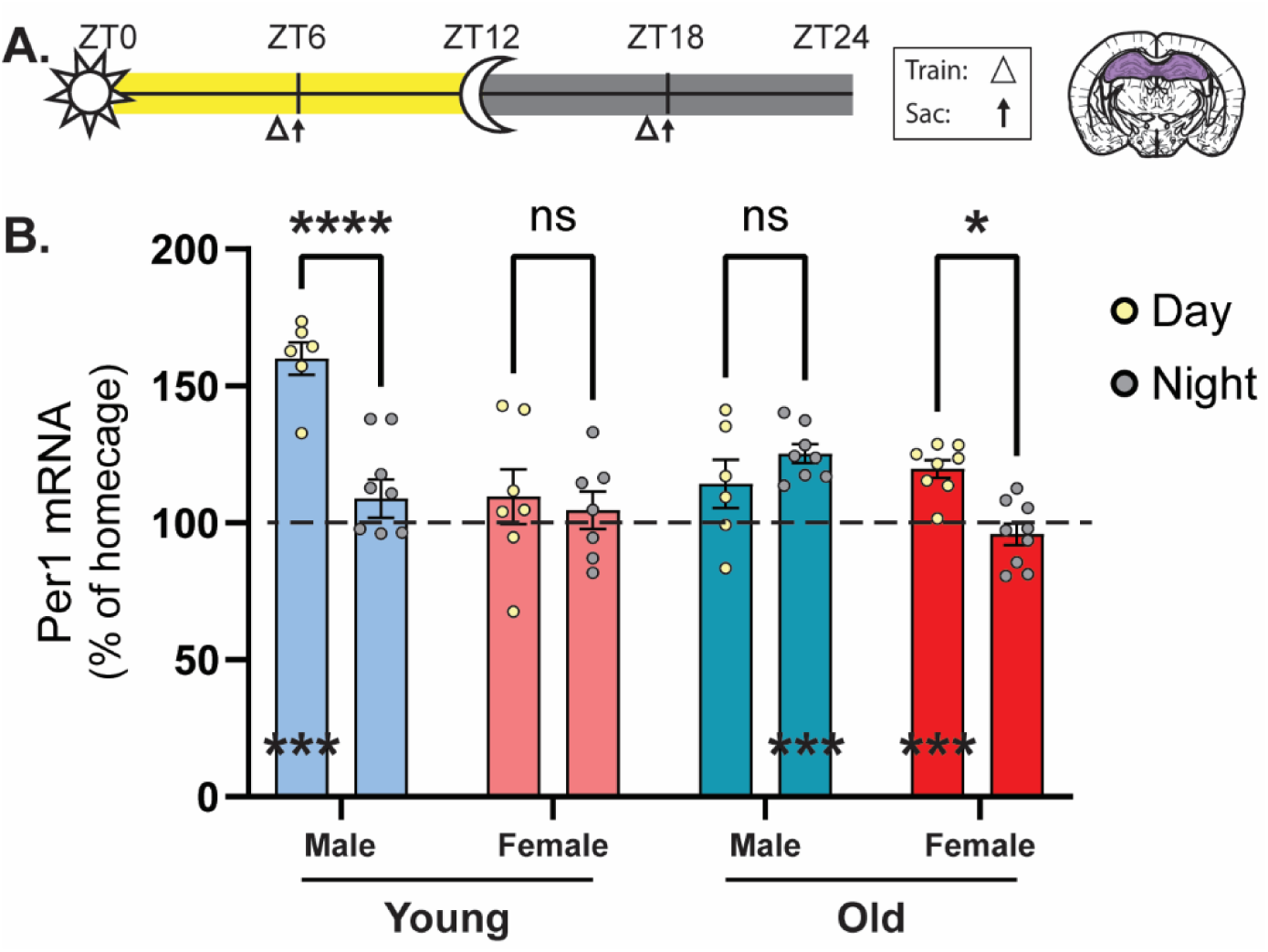
*Per1* is induced by learning, generally in tandem with memory performance. **A.** Schematic for behavioral and sacrificing timepoints. Triangles indicate time of training and arrows indicate time of sacrifice. Punches were taken from the dorsal hippocampus as indicated by purple region. **B.** Young male and old female mice have more *Per1* induced by learning during the day compared to the night. Young male and old female mice have a significant induction of *Per1* in response to learning when compared to time-matched homecage (set to 100, indicated by dotted line) during the day and old male mice have a significant induction of *Per1* during the night (n=6-9/timepoint). * = p<0.05, *** = p<0.001, **** = p<0.0001 compared to 100 or between groups. ZT = Zeitgeber Time, where ZT0 = 6am (7am DST), lights on, ZT12 = 6pm (7pm DST), lights off.

Old male mice also showed a diurnal oscillation in *Per1* expression that matched this group’s memory performance (Fig. 2D), although this pattern was opposite that of the young male/old female groups. Specifically we observed that old males showed little induction of *Per1* during the day, when memory is poor, but a significant induction of *Per1* at night when memory peaks (Fig. 3B; one sample *t-*tests comparing each group to 100, day: *t*_(5)_=1.601, p=0.1703, night: *t*_(7)_=7.407, p<0.001), although *Per1* induction was not significantly higher during the night than during the day (Sidak’s *post hoc*, *p*=0.6663). Therefore, *Per1* is roughly induced in tandem with memory in the old male mice, as well, with both memory and *Per1* induction peaking during the night.

Finally, in the young females, we observed no change in *Per1* induction between the day and night, consistent with their lack of diurnal oscillations in memory performance (Sidak’s *post hoc*, *p*=0.9721). Surprisingly, despite showing robust memory at both timepoints, these young female mice failed to show an induction of *Per1* at either ZT5 or ZT17 (Fig. 3B; one sample *t-*tests comparing each group to 100, day: *t*_(6)_=0.9632, p=0.3726, night: *t*_(6)_=0.6702, p=0.5277). This suggests that other mechanisms beyond *Per1* must be capable of supporting this robust memory during both the day and night in young female mice.

We also looked at *Per1* levels in the homecage group to determine if age or sex alters baseline *Per1* oscillations (Suppl. Fig. 3). Under homecage conditions, we did observe a significant effect of ZT on *Per1* levels, but not an effect of age or sex (three-way ANOVA significant effect of ZT time (*F*_(1,56)_=11.72, p<0.01), ZT time x age (*F*_(1,56)_=9.996, p<0.01), no significant effect of sex, age, ZT time x sex, age x sex, or ZT time x age x sex). When comparing between day and night within each cohort we did see that young animals (both male and female) had significantly more *Per1* expression during the night compared to day, as previously reported in young males [2]. In old animals, however, there was no oscillation in homecage *Per1* in either males or females (Sidak’s *post-hocs* comparing day to night: young male: p<0.05; young female: p<0.01; old male: p>0.05; old female: p>0.05). Therefore, baseline oscillations in *Per1* seem to be impacted by age, with old mice showing blunted oscillations in homecage *Per1* levels. While interesting, these differences in baseline *Per1* do not match changes in memory performance and are therefore incapable of supporting diurnal changes in memory observed across age and sex.

No differences in total object exploration at test were observed across any comparison (Suppl. Fig. 1D; three-way ANOVA no effect of ZT time, sex, age, ZT time x sex, ZT time x age, sex x age, or ZT time x sex x age), indicating that the amount of *Per1* induced by learning was not dependent on exploration time. We did observe some habitation movement differences, however. Comparing across groups, we found that distance traveled decreased across habituation sessions for all animals, but there was a significant difference in the amount of movement across the age/sex groups (Suppl. Fig. 2G-H; mixed-effects analyses; Distance: significant main effects of day (*F*_(3.660,410)_=2.736, p<0.05) and group (*F*_(7,114)_=7.474, p<0.0001), and a significant day x group interaction (*F*_(35,560)_=1.605, p<0.05)); Velocity: significant effect of group (*F*_(7,114)_=7.220, p<0.0001), but no significant effect of habituation day or day x group interaction). Post-hoc tests comparing distance or speed within each session showed that female mice, regardless of age, typically showed more activity than the male mice, especially on days one and six. In general, regardless of the time of day, female mice showed slightly more movement across habituation than male groups, with old females showing the most movement.

Overall, we found that learning-induced *Per1* expression patterns largely mirror diurnal memory performance patterns across age and sex, with larger *Per1* induction corresponding to better memory performance in young males, old males, and old females. Young females show steadily robust memory performance across the diurnal cycle that is accompanied by a consistent lack of *Per1* induction, suggesting some other mechanism beyond *Per1* must be capable of supporting spatial memory in these young female mice.

### Circadian free-running periods are similar across both age and sex

Our results indicate that age and sex impact diurnal patterns of memory performance in unexpected ways. It is possible that some of these alterations are driven by age- and sex-related changes in the animals’ circadian activity patterns. There are very few studies that assess how sex impacts the free-running period, sleep behavior, or light pulse-induced phase resetting in mice [22,23]. Thus, we next assessed these circadian behaviors in each group of animals.

In this experiment, we first monitored the activity of well-entrained mice for 7 days under light/dark (LD) conditions before releasing the mice into constant darkness (DD). During the DD period, all zeitgebers were removed, allowing the animals’ endogenous circadian rhythm to drive sleep/wake patterns. After 10 days in constant darkness, we assessed the ability of each group of mice to show photic phase resetting by exposing them to a 50-lux light pulse for 30 minutes beginning at CT16.75. Of note, similar light pulse protocols typically drive a substantial phase delay in young male mice but little to no phase shift in old male mice [18,24]. Following light pulse exposure, all mice remained in constant darkness for an additional 10 days to identify any phase shifts (Fig. 4A). Across the entire experiment, the activity pattern of each animal was constantly assessed with an infrared activity monitor mounted at the top of the cage.We first measured the circadian period (free-running tau, τ) under each lighting condition: LD, pre-pulse DD, and post-pulse DD to determine how age and sex affect the animals’ circadian activity patterns. We found that free-running τ changed across the three experimental phases but neither age nor sex had an impact (Fig. 4B; three-way ANOVA, significant effect of lighting conditions (*F*_(1.902,55.17)_=29.50, p<0.0001) but no effect of age, sex, lighting condition x age, lighting condition x sex, or lighting condition x age x sex). Post-hoc tests revealed that the young male and young female mice had significantly longer free-running periods during the LD phase compared to the pre-pulse DD phase (Sidak’s *post-hoc*, young male: p<0.05, young female: p<0.05), but this change between LD and DD was not significant for old male (p=0.9979) or old female (p=0.2667) mice. This is consistent with previous reports that, in the absence of zeitgebers, C57 mice show a free-running period of less than 24h and that the free-running period tends to lengthen as mice age [11,25]. Following the light pulse, there was no significant difference between the LD τ and the post-pulse DD τ or the pre-pulse and post-pulse period length. No other comparisons showed a difference in free-running τ across groups.

**Figure 4.**
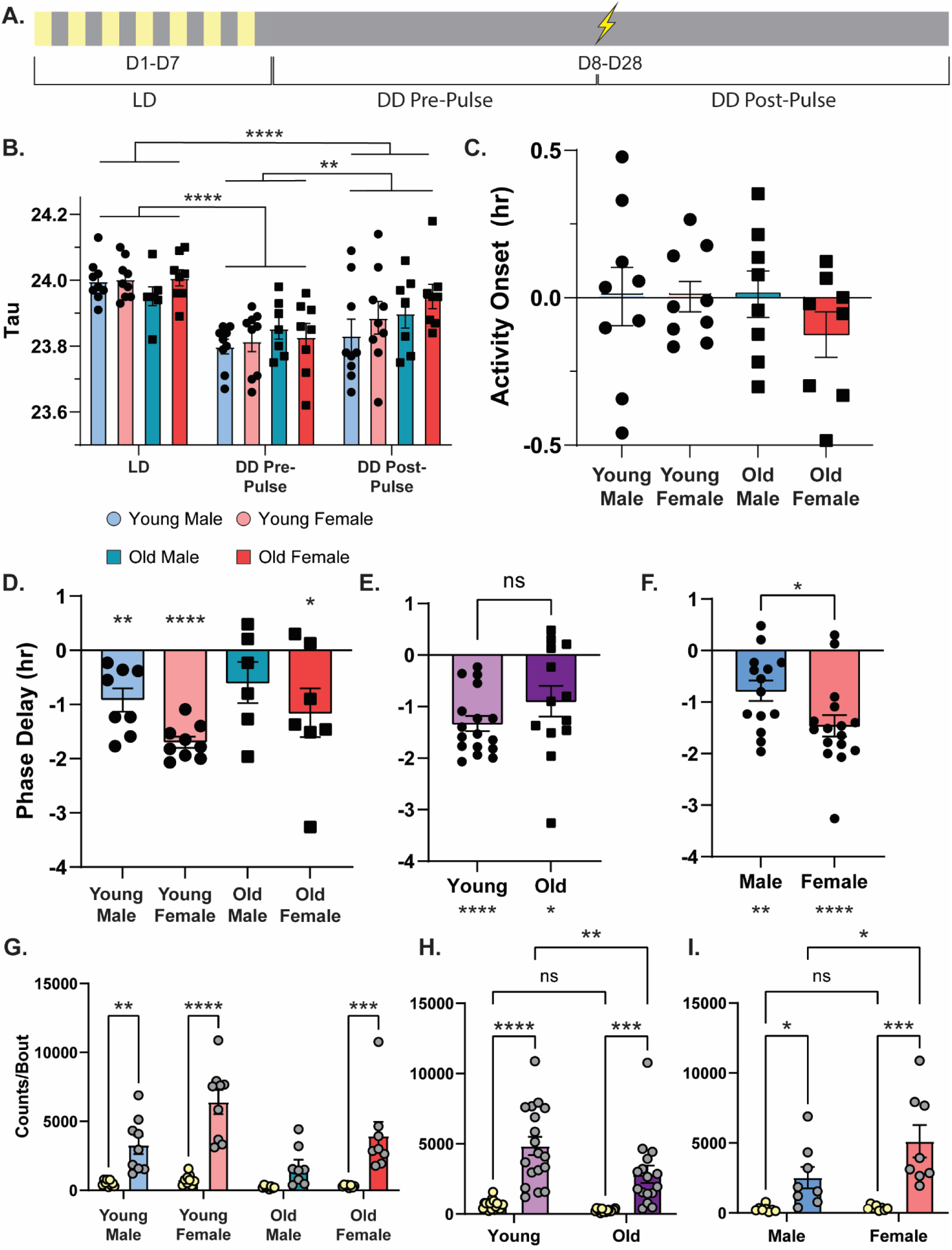
Circadian period is independent of age/sex, old males show deficits in light response and activity. **A.** Schematic for activity monitoring timeline with light pulse at CT17 represented as a lightning bolt. **B.** Circadian period under 12h light/12h dark (LD), dark/dark (DD) pre-pulse, and DD post-pulse in young male (light blue with circles), young females (light red with circles), old males (dark blue with squares), and old females (dark red with squares) (n=7-9/cohort). **C.** Time of activity onset in hours under light/dark (LD) conditions with 0 indicating time of lights turning on comparing young male, young females, old males, and old females (n=8-9/cohort). **D.** Phase delay following light pulse at CT17. Young female mice had robust phase delay. Young male and old female mice had moderate phase delay. Old male mice had minor phase delay (n=6-9/cohort). **E.** Phase delay by age comparing young mice (light purple with circles) with old mice (dark purple with squares; n=13-17/cohort). **F.** Phase delay by sex comparing male mice (blue) with female mice (red; n=14-16/cohort). **G.** The amount of activity (counts) during a bout is higher during the dark than the light in young males, young females, and old females (n=8-9/cohort). **H.** Young and old mice have more activity/bout during the dark compared to the light and young mice have more activity/bout during the dark than old mice (n=16-18/cohort). **I.** Male and female mice have more activity/bout during the dark compared to the light and female mice have more activity/bout during the dark compared to male mice (n=17/cohort). * = p<0.05, ** = p<0.01, *** = p<0.001, **** = p<0.0001 compared between groups or to 0. CT = Circadian Time.

We next collapsed all the male and female mice to determine if there is an age difference in τ length independent of sex. As before, we found an effect of phase (LD, pre-pulse DD, or post-pulse DD) on τ length but there was still no effect of age (Suppl. Fig. 4A; two-way ANOVA, significant effect of lighting conditions (*F*_(2, 62)_=30.61, p<0.0001) but no effect of age or interaction). Both ages showed longer τ during the LD than the pre- and post-pulse DD, but there were no significant differences between each age for any of the experimental phases. This suggests that age did not meaningfully impact the free-running τ across this experiment.

We also collapsed the two ages to determine if there is an effect of sex on free-running τ independent of age. Again, we found that the experimental phase had a significant impact on the free-running τ, but there was no significant impact of sex (Suppl. Fig. 4B, two-way ANOVA, significant effect of lighting conditions (*F*_(1.892, 58.66)_=30.52, p<0.0001) but no effect of sex or interaction). Again, both male and female mice showed a longer τ during the LD phase compared to the other phases but there were no significant differences between males and females in any phase of the experiment. Therefore, sex did not meaningfully impact the free-running τ across this experiment.

Together, we found that age and sex did not have a major impact on the free running circadian period, but the free-running τ did change across the experimental phases. Specifically, we consistently observed a longer τ during the LD phase compared to the pre-pulse or post-pulse DD, confirming that mice show a <24h free-running period as previously reported [11,25]. The post-pulse DD phase τ was also significantly longer than the pre-pulse DD phase (Fig. 4B; Sidak’s *post-hoc*, LD phase vs pre-pulse DD phase: p<0.0001, LD phase vs post-pulse DD phase: p<0.0001, pre- pulse DD phase vs post-pulse DD phase: p<0.01), indicating that following a light pulse the free-running τ significantly lengthens, but does not completely recover back to a LD phase.

### Time of activity onset after the lights turn off is similar across age and sex

The next measurement we looked at was activity onset, the time at which mice begin moving after the lights turn off for the day. Previous work has shown that activity onset is delayed and less precise in old male mice [11]. For this measurement, we calculated the difference between when the first bout of activity started for the active phase and the time the lights turned on averaged across the 7 day LD period (Fig. 4A). We found that all groups started activity, on average, close to when the lights turned on. Young males started moving 0.0043 hours after the lights turned on, young females 0.0040 hours after, old males 0.012 hours after, and old females 0.12 hours before the lights turned on (Fig. 4C; one-sample *t* test as compared to 0, young male: *t*_(8)_=0.04366, p=0.9662, young female: *t*_(8)_=0.07794, p=0.9398, old male: *t*_(7)_=0.1521, p=0.8834, old female: *t*_(7)_=1.621, p=0.1491). Neither age nor sex had a significant effect on activity onset (two-way ANOVA, no significant effect of age, sex, or interaction) and there was no observable age (Sidak’s *post-hoc*, male: young vs old: p>0.05, female: young vs old: p>0.05) or sex (Sidak’s *post-hoc*, young: male vs female: p>0.05, old: male vs female: p>0.05) difference for activity onset. Therefore, in contrast to previous work [11], we failed to show a significant delay in activity onset for old mice compared to young mice.

### Old male mice show impaired light pulse resetting

After 10 days in complete darkness, we exposed the mice to a 50-lux light pulse for 30 minutes (starting at CT16.75 and ending at CT17.25) to test whether each group is able to properly phase shift in response to a photic cue that initiates a phase delay in young male mice [24]. We found that there was a significant effect of sex, but not age on phase resetting to the light pulse (two-way ANOVA, significant effect of sex (*F*_(1,26)_=5.264, p<0.05), no effect of age or interaction), with the female mice showing a larger phase delay than the males. As expected, young male mice showed a moderate phase delay with this stimulation, 0.9175 hours, significantly different from zero (Fig. 4D; one-sample *t* test, young male: *t*_(7)_=4.273, p<0.01). Young and old female mice also showed successful phase shifting, with a phase delay, of 1.698 hours and 1.153 hours, respectively, both significantly different from zero (Young: *t*_(8)_=16.12, p<0.0001; Old: *t*_(6)_=2.558, p<0.05). Only the old male mice failed to show a significant phase delay, only shifting 0.5850 hours, not significantly different from zero (*t*_(5)_=1.572, p=0.1768). This pattern is consistent with past findings that young, but not old male mice show effective phase delays with similar photic stimulation [24] and extends these results to demonstrate that both young and old female mice show significant phase delays with the same stimulation.

We also collapsed across age and sex to better understand the impacts of each variable on photic resetting. We first collapsed across sex to more directly compare young and old mice. Here, we observed no significant difference in photic resetting across the two ages (Fig. 4E; unpaired *t* test, *t*_(28)_=1.405, p=0.1710), with both groups showing significant phase shifts compared to zero (young mice: *t*_(16)_=8.990, p<0.0001, old mice: *t*_(12)_=3.007, p<.05). Although the groups were not significantly different, young mice did show a stronger phase delay (1.331 hours) compared to old mice (0.8954 hours), suggesting that old age modestly dampens photic phase resetting.

Next, we collapsed the data across age to better compare the effect of sex. Here, we found that female mice had a significantly larger phase delay than male mice (Fig. 4F; males: 0.7793h; females: 1.459h; unpaired *t* test, *t*_(28)_=2.335, p<0.05), although both groups showed a significant phase delay compared to zero (one sample *t*-tests: male mice: *t*_(13)_=3.908, p<0.01, female mice: *t*_(15)_=6.978, p<0.0001). This suggests that female mice show a stronger photic resetting response than males, although both groups are capable of responding appropriately to the light pulse.

Overall, we found that photic phase delays are both sex- and age-dependent. Female mice generally showed stronger phase resetting than males, and old mice generally showed slightly weaker phase delays, although the only group that failed to show a phase delay was our old male group. To our knowledge, this is the first time that photic resetting has been assessed in female mice of any age.

### Females are more active than males across the diurnal cycle, but aging only modestly reduces activity

In general, we observe slightly more activity in our female mice compared to male mice (e.g. Suppl. Fig. 2G-H). We therefore quantified the activity of each group during the day and night under LD conditions to determine if sex also impacted activity levels in our circadian experiment (Fig. 4). First, we looked at the counts of activity per bout, which is a measurement of how much movement the mouse had during each discrete period of activity. Here, we averaged the counts/bout during the day or night across the 7 day LD period. We found that all groups had fewer counts/bout during the light than the dark phase, with the exception of the old males, which showed no significant difference (Fig. 4G; three-way ANOVA, effect of lighting condition (*F*_(1,30)_=73.47, p<0.0001), age (*F*_(1,30)_=8.958, p<0.01), sex (*F*_(1,30)_=12.64, p<0.01), lighting condition x age (*F*_(1,30)_=4.275, p<0.05), lighting condition x sex (*F*_(1,30)_=10.06, p<0.01), no effect of age x sex or lighting condition x age x sex; Sidak’s *post-hoc*, light vs dark for: young male: p<0.01, young female: p<0.0001, old male: p=0.2730, old female: p<0.001).This suggests that all groups of animals showed more activity during the dark phase than the light phase, but this diurnal difference is dampened in old male mice.

When we collapsed across sex, we did see a significant effect of age (Fig. 4H; two-way ANOVA, effect of lighting condition (*F*_(1,32)_=58.10, p<0.0001), effect of age (*F*_(1,32)_=6.603, p<0.05), no significant interaction). Both young and old mice showed more activity per bout during the dark phase compared to the light phase (Sidak’s *post-hoc*, young light vs dark: p<0.0001, old light vs dark: p<0.001), but during the dark (active) phase, young mice were more active, with more counts/bout than old mice (Sidak’s *post-hoc*, dark young vs old: p<0.01). This suggests that young mice are more active during the dark phase than old mice.

When collapsed by age, we observed a sex effect (Fig. 4I; two-way ANOVA, effect of lighting condition (*F*_(1,32)_=70.39, p<0.0001), effect of sex (*F*_(1,32)_=10.53, p<0.01), effect of interaction (*F*_(1,32)_=9.529, p<0.01)). As before, we observed more counts/bout during the dark phase, but female mice showed more activity than males during the dark (active) phase (Sidak’s *post-hoc*, male light vs dark: p<0.001, female light vs dark: p<0.0001, dark male vs female: p<0.0001). Overall, this shows that during the active phase the young females show more movement than males, and that young mice in general have more counts/bout than the old mice.

We also looked at the length in minutes of discrete activity bouts across the diurnal cycle, and we found that sex and ZT time both influenced how active the mice were, but age did not affect activity bout length (Suppl. Fig. 4C; three-way ANOVA, effect of lighting condition (*F*_(1,30)_=67.54, p<0.0001), sex (*F*_(1,30)_=10.15, p<0.01), lighting condition x sex (*F*_(1,30)_=8.044, p<0.01); no effect of sex, lighting condition x age, age x sex, or lighting condition x age x sex). Across all of these measures, we consistently see that mice show more activity during the night than during the day, with females and young mice showing the most activity.

### Sleep duration is influenced by the sex of the mouse

We also assessed sleep behavior (sleep bout length) during each phase of the experiment (Fig. 5). Here, we identified bouts of inactivity lasting 40s or longer as a behavioral correlate of sleep [2], as previously validated with EEG [19] and we compared sleep bout length across the different groups and phases of the experiment. Focusing on the LD phase, before we disturbed the natural sleep patterns, when we compared all cohorts, we found that there was a significant difference in sleep length that depended on the ZT and the sex of the animal, but not the animal’s age (Fig. 5A; three-way ANOVA, effect of ZT (*F*_(9.6,278.4)_=101.4, p<0.0001), sex (*F*_(1,29)_=7.237, p<0.05), ZT time x age (*F*_(47,1363)_=3.679, p<0.0001), ZT time x sex (*F*_(47,1363)_=3.581, p<0.0001), no effect of age, age x sex, or ZT time x age x sex). We averaged the sleep time (in seconds) during the day and the night for each cohort and found there was a significant effect of the ZT time and of sex, but not age (Fig. 5B; three-way ANOVA, effect of ZT time (*F*_(1,29)_=805.6, p<0.0001), sex (*F*_(1,29)_=7.237, p<0.05), and ZT time x sex (*F*_(1,29)_=6.805, p<0.05), no effect of age, ZT time x age, age x sex, or ZT time x age x sex). As this suggests that sex impacts sleep more than age, we decided to collapse the data and directly compare across each variable. We first collapsed the data across sex to better determine how age impacts sleep behavior. Here, we found that while sleep length is impacted by the ZT, age does not have a significant effect on sleep length (Fig. 5C; two-way ANOVA, no effect of age (*F*_(1,31)_=1.660, p=0.2072), significant effect of ZT time (*F*_(8.429,261.3)_=93.72, p<0.0001), significant Age x ZT interaction (*F*_(47.1457)_=3.535, p<0.0001)). Interestingly, the sleep length was similar for both ages during the light phase (ZT0-ZT12) but old mice showed longer sleep bouts than young mice during theirst part of the dark phase (ZT12-ZT18) and generally less variability across the dark phase than young mice. Once again, we averaged sleep time for the day and night and found that age does not impact the average sleep time during the day or the night, although both young and old mice sleep significantly more during the night than the day (Fig. 5D; two-way ANOVA, effect of ZT time (*F*_(1,31)_=676.4, p<0.0001), no effect of age or interaction, Sidak’s *post-hoc*, young light vs dark: p<0.0001, old light vs dark: p<0.0001).

**Figure 5.**
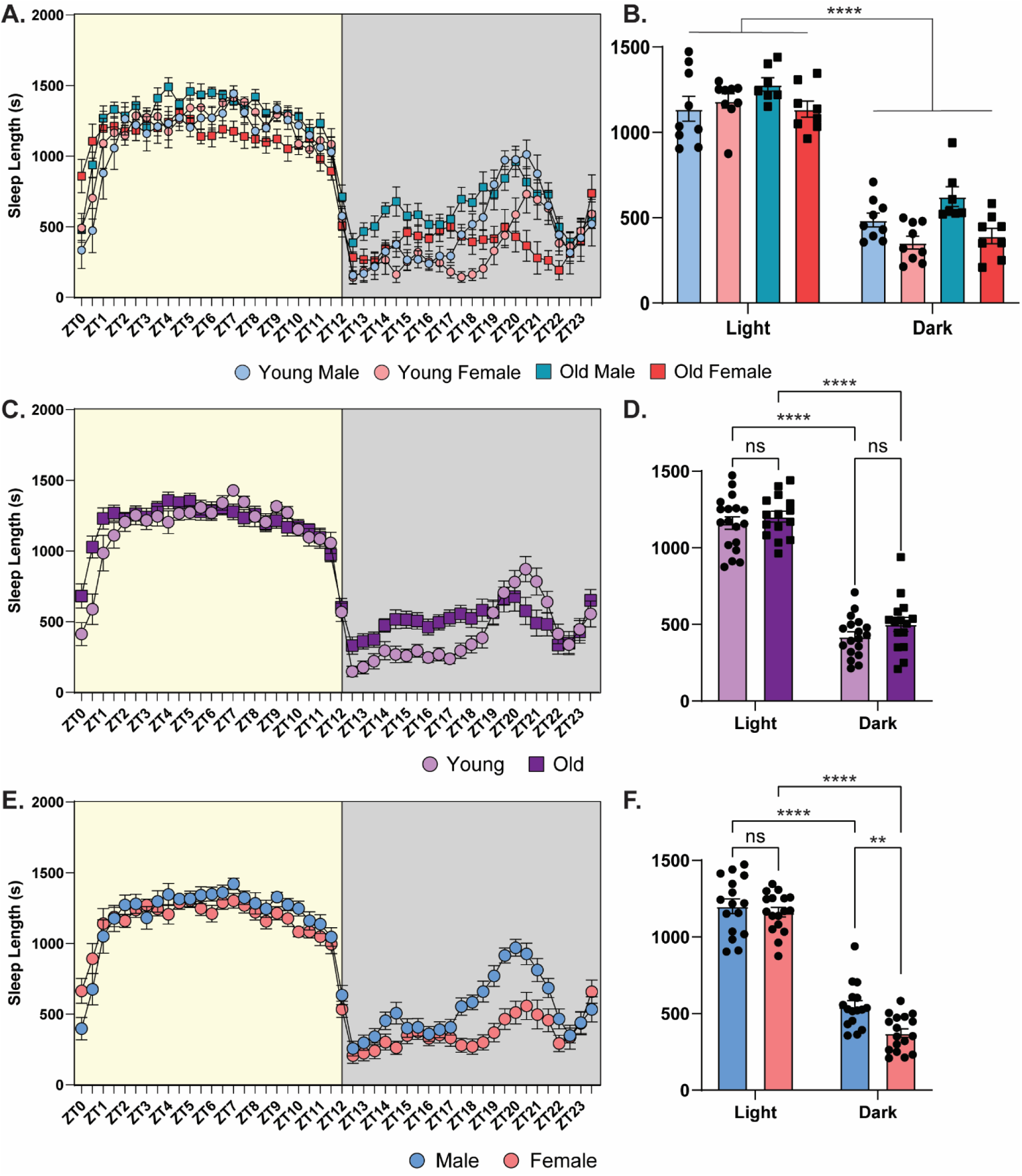
Male mice sleep more during the dark phase than female mice**. A.** Sleep length in seconds under LD conditions comparing young males (light blue with circles), young females (light red with circles), old males (dark blue with squares), and old females (dark red with squares; n=7-9/cohort). **B.** Average sleep length in seconds during the light phase and the dark phase for young males, young females, old males, and old females. All groups slept more during the light phase than dark phase (n=7-9/cohort). **C.** Sleep length in seconds under LD conditions comparing young (light purple with circles) and old mice (dark purple with squares; n=15-18/cohort). **D.** Average sleep length in seconds during the light phase and the dark phase for young and old mice. All groups slept more during the light phase than dark phase (n=15-18/cohort). **E.** Sleep length in seconds under LD conditions comparing males (blue) and females (red; n=16-17/cohort). **F.** Average sleep length in seconds during the light phase and the dark phase for males and females. All groups slept more during the light phase than dark phase and the males slept more during the dark phase than the females (n=16-17/cohort). ns = not significant ** = p<0.01, **** = p<0.0001 compared between groups. ZT = Zeitgeber Time, where ZT0 = 6am (7am DST), lights on, ZT12 = 6pm (7pm DST), lights off.

Next, we tested the impact of sex on sleep behavior, collapsing across the two ages (Fig. 5E-F). We found that sleep length is influenced by sex, with similar sleep lengths observed during the inactive phase (ZT0-12) and early active phase (ZT12-ZT17), but with females showing shorter sleep bouts than males in the late active phase (ZT17-ZT22) (Fig. 5E; two-way ANOVA, significant main effect of ZT (*F*_(9.025, 279.8)_=95.21, p<0.0001), sex (*F*_(1,31)_=5.658, p<0.05), and ZT x Sex interaction (*F*_(47, 1457)_=3.427, p<0.0001)). When we looked at the average sleep duration during the day and night we once again found that sex had a significant effect. While both sexes slept more during the day than the night and sleep lengths were similar during the day, males showed more sleep at night than females (Fig. 5F; two-way ANOVA, significant main effect of ZT time (*F*_(1,31)_=828.2, p<0.0001), sex (*F*_(1,31)_=5.658, p<0.05), and ZT time x sex interaction (*F*_(1,31)_=7.285, p<0.01), Sidak’s *post-hoc*, dark male vs female: p<0.01, male light vs dark: p<0.0001, female light vs dark: p<0.0001). Overall, sex significantly impacts sleep length in mice across the diurnal cycle, especially during the dark phase, while age does not. In general, female and male mice, regardless of age, sleep similar amounts during the day, but at night males will sleep more than females indicating that females are more active than males. To our knowledge, this is the first work to investigate how both age and sex impact sleep patterns in mice.

## Discussion

Our results show that diurnal oscillations in memory depend on both age and sex.

Our previous work showed that young male mice learned best during the day and worst at night [2] and that old male mice show memory impairments during the daytime [4].

Here, we extend this work to show that: 1) old male mice unexpectedly show better spatial memory at night than during the day, 2) young female mice show robust memory across the diurnal cycle and are resilient to time-of-day impairments in memory, and 3) old female mice show an emergence of diurnal memory oscillations, with better memory during the daytime. We also found that, in general, learning-induced *Per1* levels matched memory performance, consistent with our hypothesis that *Per1* exerts diurnal control over long-term memory consolidation. Indeed, our group [2–4,17] and others [26,27] have shown that *Per1* plays a causative role in long-term memory formation, consistent with the idea that *Per1* can regulate memory across the diurnal cycle. One important exception was the young female group, which showed robust memory during both the day and the night despite showing no learning-induced increases in *Per1*. This suggests that females may rely on alternate mechanisms to support memory in young adulthood, but as these animals age, diurnal oscillations in both memory and *Per1* emerge (discussed below). Finally, we assessed the circadian rhythms, sleep behavior, and photic phase resetting in each group and found that circadian rhythms and sleep were similar across groups (with slightly more sleep observed in males and old animals), but light pulse phase delays were stronger in females than males, an effect that was dampened with age in both sexes. Therefore, while circadian activity patterns were not grossly different across age or sex, old and male mice may show a reduced ability to rapidly re-entrain their circadian rhythm to fluctuations in the day/night cycle. Nonetheless, as our memory tasks were conducted under a stable 12h light/dark cycle, with prominent zeitgebers, it is unlikely that these slight alterations in the circadian rhythm drove our effects on memory.

Our most striking and novel finding was that young female mice showed no impact of time-of-day on long-term spatial memory performance, an effect that occurred despite the lack of *Per1* induction during either the day or night. As these female mice age, however, diurnal fluctuations in both memory and *Per1* induction emerge that match the oscillations observed in young males. This suggests that females are not susceptible to diurnal ebbs in memory performance and must rely on a mechanism outside of *Per1* to support this robust performance, as hippocampal *Per1* inductions are absent. Although the mechanism for this effect is unknown, it is tempting to speculate that estrogens (especially 17β-estradiol, E2) play a key role. E2 is known to improve long-term memory [28–30] and high levels of estradiol in young female mice might be capable of supporting robust memory across the diurnal cycle despite the lack of *Per1* induction.

Further, as estradiol circulation drops precipitously in old female mice [31], this reduction in estradiol could explain why old females show a male-like emergence of diurnal oscillations in both memory and *Per1*; this removal of estradiol could re-engage the machinery that normally exerts diurnal control in male mice. Unfortunately, we did not confirm estrous phase in these experiments, as we did not predict this result and wanted to avoid potential stress effects in females due to vaginal cytology [32,33].

Nonetheless, it is clear that young female mice show consistent memory performance across the diurnal cycle, even when collapsed across estrous phase, and this changes in old age, when female mice experience reproductive senescence.

Our old female mice were between the ages of 19-22 months at the time of training, meaning they were no longer cycling [31]. Although we did not confirm this in our current cohort, our lab has reliably shown acyclicity in 19-month-old female rodents via vaginal cytology (Suppl. Fig. 5). Acyclic mice show either a complete lack of cell type within the vaginal smear (Suppl. Fig. 5A) or the presence of a persistent diestrus state (Suppl. Fig. 5B). Persistent diestrus in acyclic mice is characterized by an inability to move from the diestrus phase but is distinguishable from cyclic estrus in younger females (Suppl. Fig. 5C) by the amount of leukocytes present within the smear (far less in acyclic mice). In female rodents, estradiol levels have been observed to drop by 50% at 13 months of age, and 75% by 17 months [31], indicating that the 19-22-m.o. female mice used here likely had a severe reduction in estradiol at the time they were trained and tested. Together, this is consistent with the hypothesis that old females experience a drop in estradiol that enables *Per1* to emerge as a diurnal regulator of memory, something that will need to be tested in future studies.

There are a few papers that look at both aging and sex as factors to the learning process [34–36] or circadian rhythm [22,23] in rodents. The existing literature on circadian rhythms and aging has not analyzed the data using sex as a factor for majority of the analyses, although one study did show that the free-running τ was not significantly different between young male and female mice of the same strains [23], consistent with our findings here. As none of these previous papers looked at photic resetting, there is little known about sex differences in phase advances or delays in rodents. Here, we present the first comprehensive evidence for sex- and age-mediated differences in diurnal memory, *Per1* learning-induced changes, and light pulse responding.

Another interesting and unexpected finding in this study was our discovery that old male mice do not simply show weak memory across the diurnal cycle but instead show a drastic shift in when memory is best and worst relative to their young counterparts.

Previous work has consistently shown that old male mice show impaired memory in non-aversive spatial tasks [3,14], but the literature on age-related memory impairments typically only looks at a single timepoint across the diurnal cycle, usually during the day. Here, we found that old male mice showed better memory performance at night than during the day. While we found that old males showed no significant differences in learning-induced *Per1* between the day and the night, there was a trend toward a larger induction in *Per1* at night that corresponded to better memory, consistent with our overarching hypothesis that *Per1* regulates the strength of long-term memory consolidation across the diurnal cycle. Why *Per1* and spatial memory peak at a different time of day in old male mice compared to young males is unclear, however, something that should be addressed in future studies.

Finally, we also looked at many different circadian rhythm measures to assess how the circadian rhythm differs between males and females, and how it changes during the aging process. As the majority of the work to date looking at circadian rhythms and aging has used only male rodents [10,24,37,38], this is a critical piece of information needed to understand the complex relationship between aging, sex, and diurnal regulation of memory. We hypothesized that we would see the greatest differences between the young and old mice regardless of sex, but we actually found that there were more drastic differences between the sexes than across the different ages. One key finding was that both young and old mice showed fairly normal circadian activity patterns, but general activity levels and the amount of movement per active bout both decreased in the old males but not the old females. Additionally, sleep was slightly elevated during the active phase for males (both young and old), with old age having surprisingly little effect on sleep behavior. The group with the least sleep during the active phase was the young females, also the group to show the most efficient photic phase delay in the light pulse experiment. Finally, we found that old animals show a slight deficit in photic phase resetting that may make them less adaptable to fluctuations in the circadian cycle, but this likely had no effect on the memory task described here, as all memory tasks were conducted under a 12h light-dark cycle, when all animals showed similar ∼24h period lengths.

Overall, we found that there are sex and age differences regarding the best time of day for memory performance. In addition, the mechanisms driving the time-of-day effect may change across the lifespan and be different between the sexes. Here, we showed that there is an intrinsic link between the circadian rhythm and memory in male and female mice and both processes change as mice age. Finally, this work provides additional evidence that the clock gene *Per1* may be capable of regulating diurnal oscillations in spatial memory within the dorsal hippocampus. Changes in *Per1* across the lifespan may therefore contribute to age-related changes in memory in male and female mice.

## Data Availability

The data presented here are available upon reasonable request from the corresponding author.

## Ethics Declaration

All experiments were performed according to US National Institutes of Health guidelines for animal care and use and were approved by the Institutional Animal Care and Use Committee of the Pennsylvania State University.

## Consent for Publication

All authors read and approved the final manuscript for publication.

## Author Contributions

Study concept and design: LB, GCP, and JLK. Acquisition of data: LB, GCP, MJV, CWS, ARB. Analysis of data: LB, GCP, MJV, ARB, AP, MJJ. Drafting of manuscript: LB, GCP, and JLK. Final approval of manuscript: All authors.

## Funding

This research was funded by NIH grants R01AG074041 (JLK), K99/R00AG056586 (JLK) and R21AG068444 (JLK), McKnight Brain Research Foundation/AFAR Innovator Award in Cognitive Aging and Memory Loss (JLK), American Federation for Aging Research Grant #A21105 (JLK), startup funds from the Eberly College of Science and Department of Biology at Pennsylvania State University (JLK) and the National Institute on Aging under Grant T32 AG049676 to The Pennsylvania State University (LB & CWS).

**Abbreviations**

BMAL: brain and muscle ARNT-like

CLOCK: circadian locomotor output cycles kaput

*Cry*: *Cryptochrome*

CT: circadian time

DD: dark/dark phase

DH: dorsal hippocampus

DI: discrimination index

E2: 17β-estradiol

LD: light/dark phase

OLM: object location memory

*Per1*: *Period1*

SCN: suprachiasmatic nucleus

ZT: zeitgeber time

