## Supplementals for "Age and Sex Influence Diurnal Memory Oscillations, Circadian Rhythmicity, and *Per1* Expression"

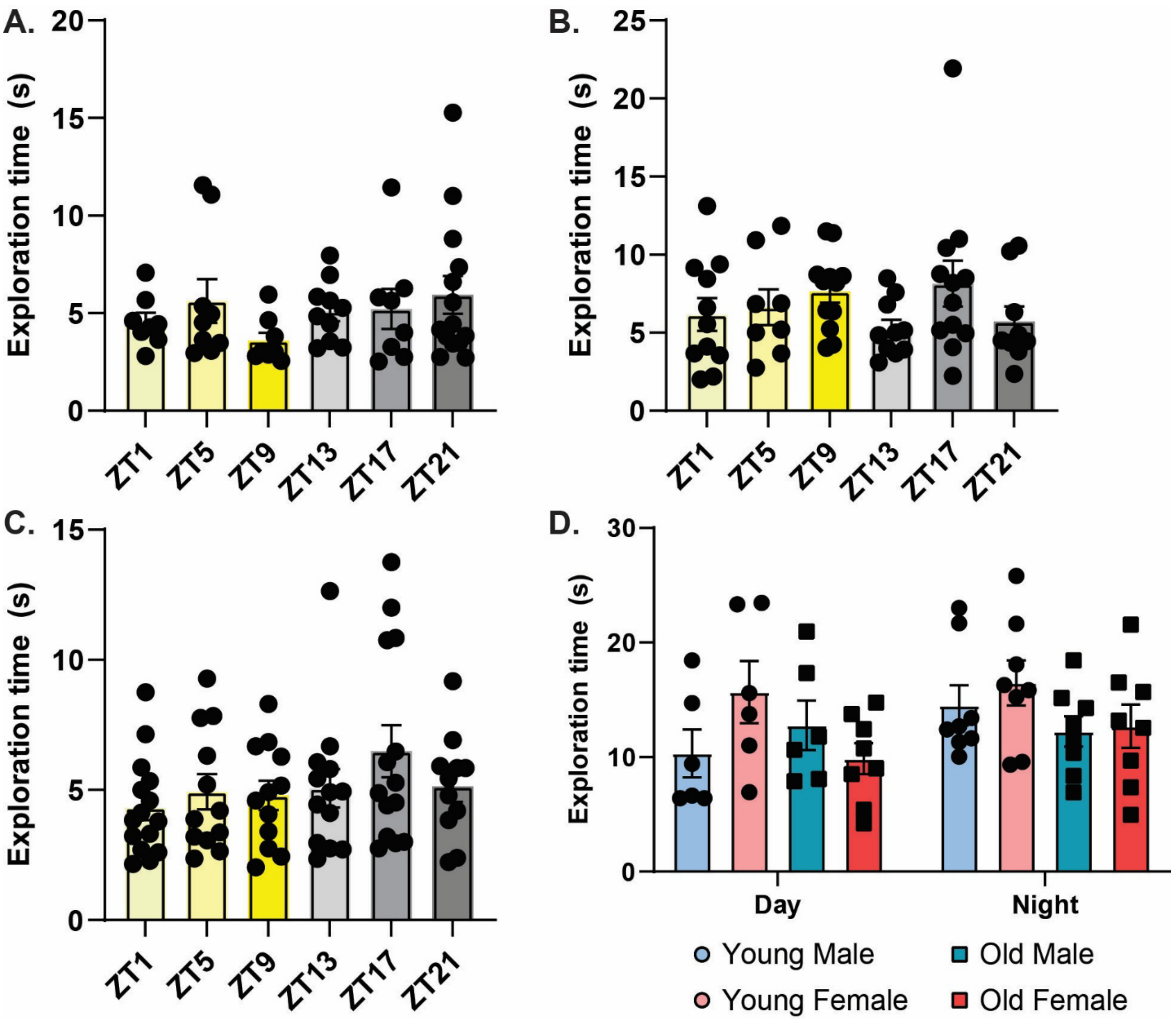


**Supplemental Figure 1.** Total exploration does not differ across the diurnal cycle. **A.** Total object exploration during the test session for the young female circadian memory experiment (Fig. 1C-D; n=8-14/timepoint). **B.** Total object exploration during the test session for the old female circadian memory experiment (Fig. 1E-F; n=8-12/timepoint). **C.** Total object exploration during the test session for the old male circadian memory experiment (Fig. 2; n=11-15/timepoint). **D.** Total object exploration during the training session for the *Per1* induction experiment Fig. 3; 6-8/timepoint). ZT = Zeitgeber Time, where ZT0 = 6am (7am DST), lights on, ZT12 = 6pm (7pm DST), lights off.

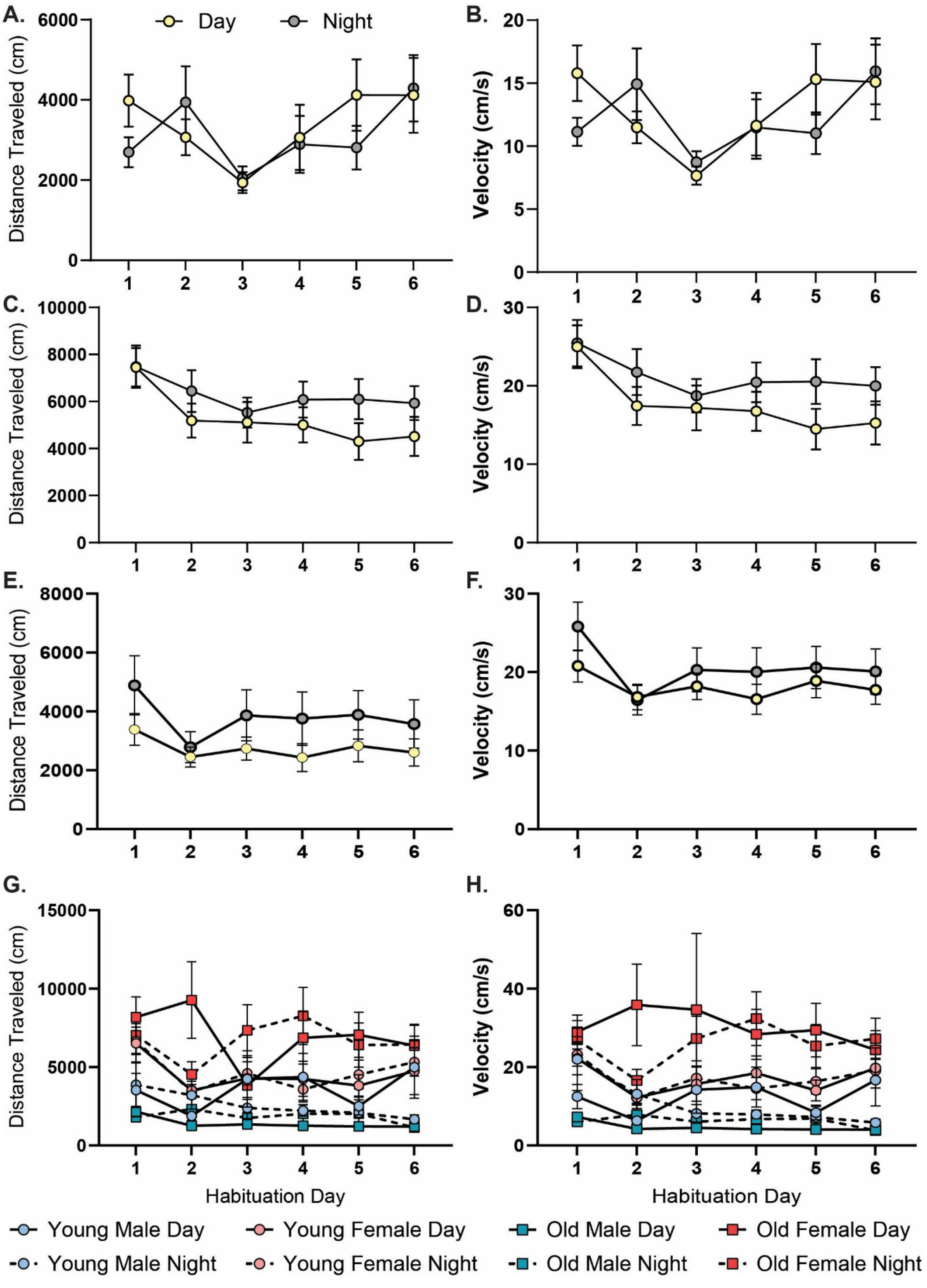

A.
B.
C.
D.
E.
F.
G.
H.
Day
Night
Distance Traveled (cm)
Velocity (cm/s)
Habituation Day
Young Male Day
Young Female Day
Old Male Day
Old Female Day
Young Male Night
Young Female Night
Old Male Night
Old Female Night

**Supplemental Figure 2.** Mouse distance traveled and velocity is similar across the diurnal cycle, with no effect of age but a sex effect, with females showing more movement during habituation. **A.** Distance traveled (cm) and **B.** velocity (cm/s) of young female mice habituated during the day (yellow symbols; ZT1, ZT5, ZT9) and at night (grey symbols; ZT13, ZT17, ZT21) across the 6 habituation days from the young female circadian experiment (Fig. 1C-D; n=25-32/timepoint). **C.** Distance traveled (cm) and **D.** velocity (cm/s) of old female mice habituated during the day and at night across the 6 habituation days from the old female circadian experiment (Fig. 1E-F; n=31/timepoint). **E.** Distance traveled (cm) and **F.** velocity (cm/s) of old male mice habituated during the day and at night across the 6 habituation days from the old male circadian experiment (Fig. 2; n=38-39/timepoint). **G.** Distance traveled (cm) and **H.** velocity (cm/s) across the 6 habituation days from the *Per1* induction experiment (Fig. 3; n=12-17/cohort).

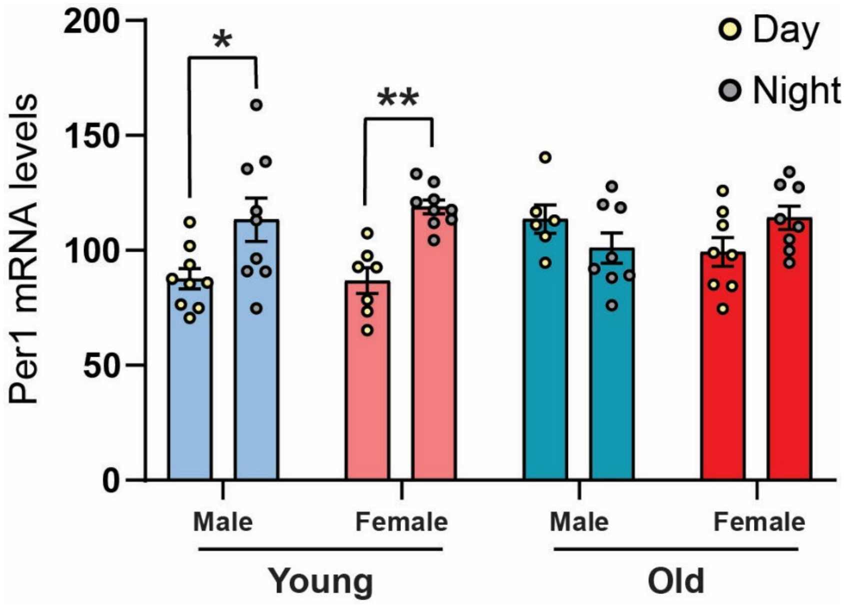


**Supplemental Figure 3.** *Per1* mRNA levels in homecage young male (n=9/cohort) and female (n=7-9/cohort) mice were significantly higher during the night compared to the day, but no differences were seen in old male (n=6-8/cohort) and female (n=8/cohort) mice. * = p<0.05, ** = p<0.01.

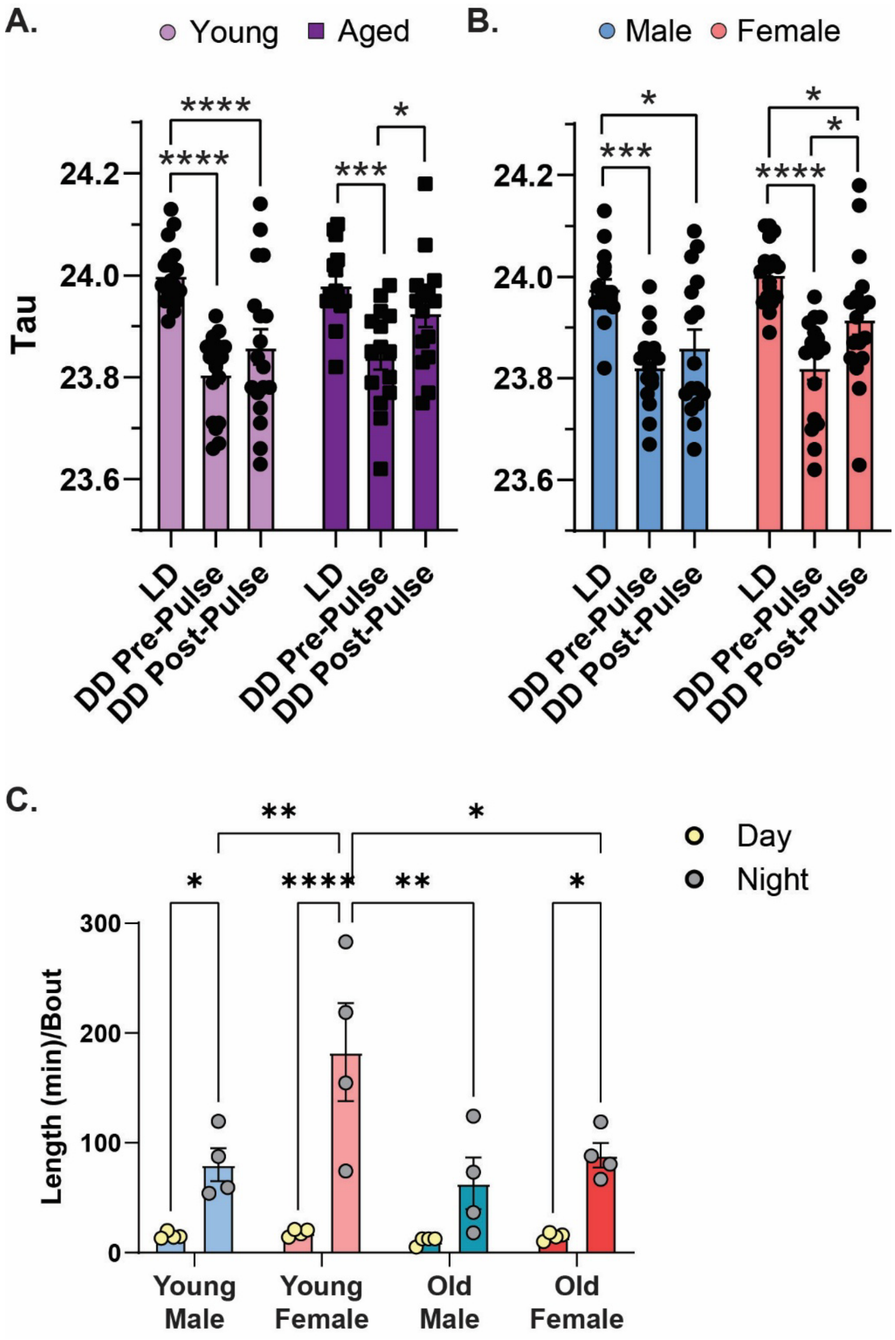


**Supplemental Figure 4.** Circadian period and activity as compared by sex and age. **A.** There is no effect of age on the circadian period length (free-running tau; n=15-18/cohort). **B.** There is also no effect of sex on free-running tau (n=16-17/cohort) **C.** The length (in minutes) of bouts during the dark phase (grey) is significantly higher in young males, young females, and old females than the light phase (yellow) while old males show no difference (n=8-9/cohort). LD = light/dark, DD = dark/dark. ns = not significant, * = p<0.05, ** = p<0.01, *** = p<0.001, **** = p<0.0001 as compared between groups.

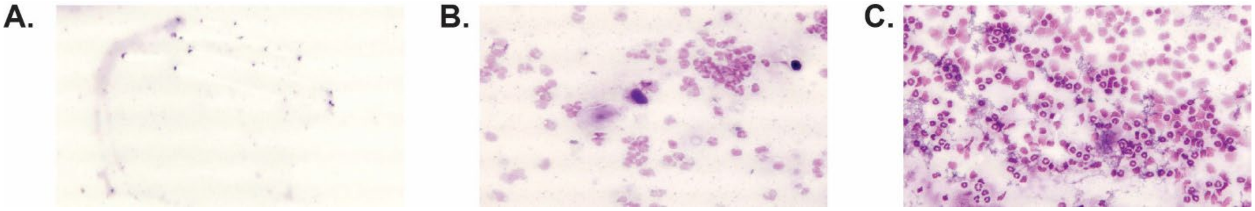


**Supplemental Figure 5.** Representative vaginal cytology smears from **A.** Acyclic 19-month-old mouse **B.** Acyclic Persistent Diestrus 19-month-old mouse **C.** Cyclic Diestrus 8-week-old mouse. Persistent Diestrus and Diestrus in cyclic mice is easy to differentiate due to the quantity of leukocytes present in the smear (smaller quantity for persistent diestrus).
